# Using arterial spin labelling to investigate spontaneous and evoked ongoing musculoskeletal pain

**DOI:** 10.1101/163196

**Authors:** Karolina A. Wartolowska, Daniel P. Bulte, Michael A. Chappell, Mark Jenkinson, Thomas W. Okell, Matthew A. Webster, Andrew J. Carr

## Abstract

Clinical pain is difficult to study using standard Blood Oxygenation Level Dependent (BOLD) magnetic resonance imaging because it is often ongoing and, if evoked, it is associated with stimulus-correlated motion. Arterial spin labelling (ASL) offers an attractive alternative. This study used arm repositioning to evoke clinically-relevant musculoskeletal pain in patients with shoulder impingement syndrome. Fifty-five patients were scanned using a multi post-labelling delay pseudo-continuous ASL (pCASL) sequence, first with both arms along the body and then with the affected arm raised into a painful position. Twenty healthy volunteers were scanned as a control group. Arm repositioning resulted in increased perfusion in brain regions involved in sensory processing and movement integration, such as the contralateral primary motor and primary somatosensory cortex, mid- and posterior cingulate cortex, and, bilaterally, in the insular cortex/operculum, putamen, thalamus, midbrain and cerebellum. Perfusion in the thalamus, midbrain and cerebellum was larger in the patient group. Results of a post hoc analysis suggested that the observed perfusion changes were related to pain rather than arm repositioning. This study showed that ASL can be useful in research on clinical ongoing musculoskeletal pain but the technique is not sensitive enough to detect small differences in perfusion.

## Introduction

Some conditions cannot be easily studied using standard Blood Oxygenation Level Dependent (BOLD) functional magnetic resonance imaging (FMRI). For example, clinical pain is usually either spontaneous and ongoing or diminishes too slowly, when evoked, to be used in an event-related or block design. Musculoskeletal pain, such as shoulder pain, is particularly challenging to image using a traditional BOLD FMRI paradigm because of artefacts caused by stimulus-correlated motion. Moreover, the movement of the affected arm, so close to the head, dramatically impacts the magnetic field homogeneity of the scanner resulting in significant artefacts.

Arterial spin labelling (ASL) offers an attractive alternative to BOLD FMRI because, unlike BOLD, it does not require a clear baseline/stimulus difference and can be used to measure an ongoing state or prolonged activation. ASL has been used to measure perfusion changes related to experimental pain in healthy people (1, 2) and to clinical pain in chronic pain patients (3, 4). Pain-related perfusion has been analysed either by comparing brain perfusion during a painful and pain-free state in the patient group, comparing perfusion between the patient and control group (3, 5) or performing time-course analysis by correlating the pain ratings during imaging with the perfusion values (2, 4).

Clinically-relevant pain is of primary interest in trials of treatment efficacy (6, 7). However, it is more difficult to study than experimental pain, as it is less standardised and reproducible. Some types of clinical pain, such as after tooth extraction, can be reliably evoked in all patients (5), whereas other types are more difficult to investigate. For example, one study had to exclude seven out of 23 recruited patients because their clinical pain did not diminish quickly enough to obtain a pain-free baseline for the experimental pain session (3). Another study excluded nine out of 43 patients because they reported no pain on the day of the scan (4). Losing patients from research is a significant problem, considering how difficult it is to recruit them into longitudinal pain studies.

The aim of this study was to investigate brain perfusion during spontaneous and evoked ongoing clinical musculoskeletal pain. The hypothesis was that raising the affected arm would evoke pain in patients but not controls and that such pain would be associated with increased perfusion in pain-processing regions in patients but not controls. For the between-group analysis (patients versus controls), the hypothesis was that the difference in perfusion during the arm-down condition would reflect a spontaneous ongoing pain, while the difference in perfusion during the arm-up condition would reflect evoked ongoing pain.

## Materials and Methods

### Participants

Shoulder pain patients participating in the CSAW (Can Shoulder Arthroscopy Work, NCT01623011) trial were invited to take part in a neuroimaging study. The CSAW trial investigated the efficacy of arthroscopic decompression surgery for chronic shoulder impingement pain. The inclusion criteria for patients were subacromial pain due to tendinopathy or a partial tear, which lasted at least three months despite conservative treatment, and lack of contraindication for MRI. A detailed list of inclusion and exclusion criteria was described in the trial protocol (8).

Healthy control participants were recruited through advertisements within the Oxford hospitals. Volunteers were eligible if they were healthy, pain-free and had no contraindications for MRI.

This study was performed according to the Declaration of Helsinki and approved by the National Research Ethics Service (NRES) South Central-Oxford B Research Ethics Committee (Reference number: 12/SC/0028). All participants gave written informed consent to participate.

### Experimental design

Two ASL datasets were collected for each subject. Firstly, patients were scanned while in a supine position with their arms along the body (arm-down condition) (Figure 1). This scan measured cerebral perfusion during a pain-free state or spontaneous ongoing pain (depending on whether patients experienced spontaneous ongoing pain at rest). During the subsequent scan, patients were asked to move the affected arm into a position that evoked ongoing clinical pain (arm-up condition): maximal abduction and lateral rotation within the scanner constraints. This scan measured cerebral perfusion during ongoing evoked clinical pain. As this was the last scan of the session, patients were advised to press the buzzer if the pain became intolerable, which prompted an immediate termination of the scan and the imaging session. Controls were also scanned with their arm down and up to control for the effects of arm repositioning; the side in the control group was randomised.

**Fig. 1.**
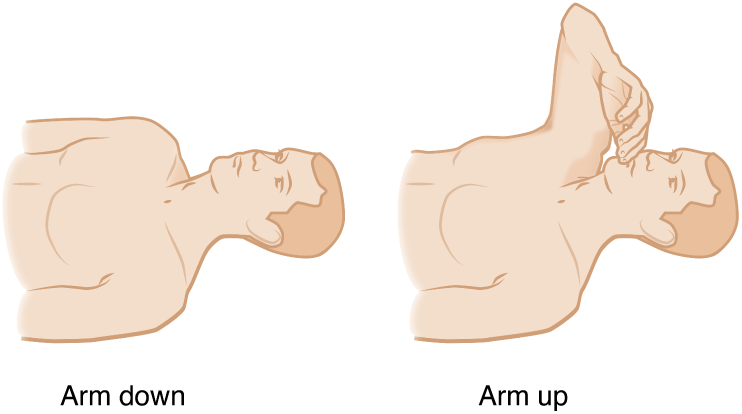
Scanning conditions. There were two scanning conditions: arm-down with the arms along the body and arm-up with the arm raised (in maximal abduction and lateral rotation within the scanner constraints).

Two ASL scans were acquired independently from each other. The arm-down scan began with a localiser, time-of-flight scan, and high-resolution structural scan. The time when the structural image was acquired was used to identify the labelling plane and to plan the ASL scan. The labelling plane was chosen based on the time-of-flight scan of the neck arteries; it was positioned perpendicularly to the axis of the vessels, at approximately the level of the second cervical vertebra (9). This was followed by shimming, the main ASL scan, two calibration scans (first using the head coil, then the body coil to correct for the coil sensitivity variations) and field map scans.

After this series of scans, patients were asked to verbally rate their average pain during the arm-down ASL scan on an 11-point numerical rating scale (NRS), with the anchors being 0 = “no pain” and 10 = “worst pain possible”. Between the arm-down and the arm-up scan, the participant was repositioned. The raised arm was padded with foam wedges for support, to prevent movement and to provide insulation from the side of the scanner. The arm-up scanning procedure consisted of re-shimming, new localizer, and time-of-flight scans, re-planning the location of the tagging plane, and a new set of ASL data. Separate calibration and field map scans were acquired for the arm-up condition to correct for potential changes due to arm repositioning. After these scans, the patient was removed from the scanner and asked to rate verbally their average pain during the arm-up scan.

### Imaging parameters

Neuroimaging was performed on a 3T Verio MRI scanner (SIEMENS, Erlangen, Germany) using a 32-channel head coil. A standard pseudo-continuous ASL sequence (pCASL) with background suppression (using pre-saturation) and an EPI readout was used (9, 10). The sequence parameters were: repetition time (TR) 4000ms, echo time (TE) 13ms, Partial Fourier = 6/8th, flip angle 90°, FOV 240 x 240mm, matrix 64 x 64. There were 24 slices collected in ascending order. The in-plane resolution was 3.75 x 3.75mm^2^, and the slice thickness was 4.5mm with a 0.5mm gap. There were six postlabelling delays, 250ms apart (250, 500, 750, 1000, 1250 and 1500ms) with label duration 1400ms and extra post-labelling delay for superior slices of 45.2ms each. These parameters were validated for modelling of kinetic curves across the whole brain in human subjects (9), including pain studies (2). There were 120 volumes (10 epochs of six label-control pairs) collected in just over eight minutes.

Two calibration images were acquired without background suppression or labelling, one using the head coil and the other using the body coil for signal reception. The TR was 6000ms and all the other parameters were the same as for the main pCASL sequence. Field maps were collected with the same orientation and voxel size as the pCASL and TR = 400ms, TE1 = 5.19ms, TE2 = 7.65ms, flip angle 60°. A structural image was acquired using the 3D MPRAGE sequence with the following parameters: TR = 2040ms, TE = 4.7ms, TI = 900ms, flip angle *8*°, FOV read 192mm, 1mm isometric voxels.

### Analysis pipeline

The data were analysed using the Oxford Centre for Functional Magnetic Resonance Imaging of the Brain (FMRIB) Software Library (FSL) tools version 5.0.9 (http://fsl.fmrib.ox.ac.uk) and Matlab (Mathworks, Natick, MA, USA). An overview of the multi-stage analysis pipeline is presented in Figure 2 and the scripts are available on GitHub (https://github.com/ka-wa/MRIscripts).

**Fig. 2.**
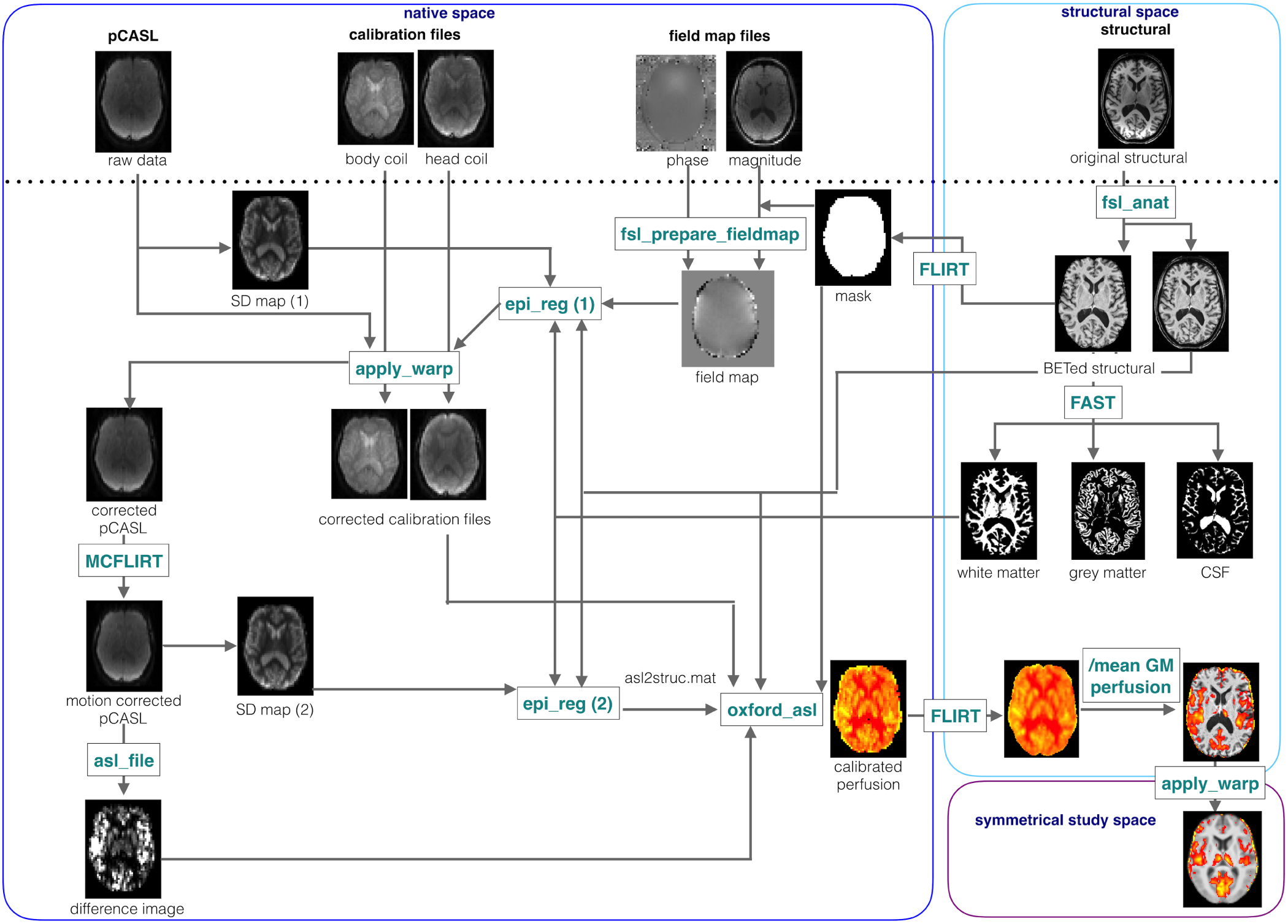
Analysis pipeline. A schematic presentation of pre-processing and processing steps used to generate perfusion maps and to transform them, first, from the native ASL space to each subject’s structural space and then to the symmetrical study-specific template.

As standard brain-extraction does not work well on ASL data, a brain-extracted structural image was used to mask non-brain tissue. This image was generated using the fsl_anat (http://fsl.fmrib.ox.ac.uk/fsl/fslwiki/fsl_anat) and transformed to the ASL space using the FLIRT registration (http://fsl.fmrib.ox.ac.uk/fsl/fslwiki/FLIRT). Fsl_anat was chosen over BET skull-stripping (11) because it corrects the structural data for RF field inhomogeneity, performs FAST segmentation (12) and removes more non-brain tissue, which was useful for masking.

The same mask was used to remove non-brain tissue from the field magnitude image, which was then used in fsl_prepare_fieldmap to create a field map image. The field map image was used as an input by the epi_reg (http://fsl.fmrib.ox.ac.uk/fsl/fslwiki/FLIRT/UserGuide*#*epi_reg) together with a structural and a brain-extracted structural image, a white matter mask from FAST segmentation and a variance map of the ASL data. The variance map was chosen as the ASL space reference image because it provided a better between-tissue contrast than ASL data or calibration files. The key step, which improved the registration, was to use epi_reg to correct the raw ASL data and calibration files for distortions due to B0 inhomogeneities and to create a good transform between the native ASL space and the structural space.

The adjusted ASL data were motion-corrected using MCFLIRT (13). Then, the label-control pairs were subtracted and the result averaged at each post-label delay using asl_file, which is a part of the BASIL (Bayesian Inference for Arterial Spin Labelling MRI) toolbox (14). A second variance map was created from the corrected ASL data to be used by epi_reg to generate the final transform between the ASL and the structural space. Perfusion was quantified using the oxford_asl script from the BASIL toolbox with the following inputs: multi post-label delay label-control difference image, brain-extracted structural image, transformation between the native and the structural space, and between the structural and the symmetrical study space (see below), variance map as a reference image, spatial priors and inversion efficiency of 0.88 (9). The voxel-wise calibration option was used to correct for a possible change in coil sensitivity due to arm repositioning (15).

The individual perfusion images were transformed to a study-specific template using an affine (FLIRT) and then nonlinear (FNIRT) registration, like in a standard FEAT analysis, and smoothed with a sigma = 3mm (FWHM = 7.05 mm) Gaussian kernel before the group analysis. The study-specific template was created by averaging structural scans of an even number of patients and controls, affine transforming them to the standard MNI152_T1 template, creating a mean and an x-axis flipped mean image, then averaging them to create a new template. Following this, the individual structural scans were registered to the template using an affine and non-linear transformation, averaged, flipped along the x-axis and averaged again to create a study-specific symmetrical template.

Optimised registration, resulted in an improved registration in comparison to FLIRT or epi_reg, with calibration images used as a reference. The mean cost function across all subjects for standard registration of a calibration image to structural image, using FLIRT, was 0.20 (95% CI 0.18 to 0.22). The mean cost function for registration of calibration image to the structural image, using epi_reg with field map correction, was 0.17 (95% CI 0.16 to 0.19). Finally, the mean cost function for the optimised registration, using epi_reg with variance map as an input, was the lowest: 0.16 (95% CI 0.15 to 0.18). The cost function for the two latter approaches was significantly better than it was for FLIRT.

For patients with affected left shoulders and controls with the stimulation on the left shoulder, data were flipped along the x-axis so that the activation was consistently ipsi- or contralateral to the stimulation.

### Statistical analysis

Baseline characteristics in the patient and control group were compared using two-sample t-tests, and the Wilcoxon signed-rank test for continuous variables and Fisher’s exact tests for categorical variables.

A regression analysis was used to analyse differences in mean grey matter perfusion and arterial transit time (ATT) between patients and controls, and between arm-down and arm-up condition, as well as interactions between those two factors, controlling for age, sex and clustering within each subject. Statistical analysis of non-imaging data was performed using STATA version 12.1 (StataCorp 2011).

Perfusion data were analysed using a voxel-wise analysis. The main analysis model tested the effect of condition and the interaction between the condition and group. The first explanatory variable was the condition (+1 for arm-up and −1 for arm-down), and the second explanatory variable was the product of condition and group (+1 for patients and −1 for controls); the mean grey matter perfusion, and subject-specific intercepts (+1 for each subject, 0 otherwise) were entered into the model as covariates of no interest. The variable that would normally be used to model group was not included because it would result in rank-deficiency with the variables modelling the subject-specific means. In this design, it was not possible to test the main effect of the group, as that was a between-subject factor. To investigate between-group differences, a separate model was set up with the patient group mean as the first explanatory variable, the control group mean as the second explanatory variable and mean grey matter perfusion, age, sex and subject-specific slopes (+1 for arm-up of that subject, −1 for arm-down for that subject, 0 otherwise) as covariates of no interest.

Statistical analysis of imaging data was performed using parametric methods (16, 17), corrected for family-wise error using Threshold-Free Cluster Enhancement (TFCE) and thresholded to show voxels significant at p < 0.05. The analysis was limited to voxels within the grey matter mask, common to all the datasets. Local maxima coordinates were reported after transformation from the symmetrical study-specific template to MNI space.

### Post hoc analysis

Differences in perfusion between arm-down and arm-up condition were also analysed in the patient and control group separately, using a regression model with a covariate for the group effect, subject-specific intercepts, and the mean grey matter perfusion values for each participant entered as a covariate of no interest. To investigate whether the changes between conditions in the patient group were in the same brain regions as those correlating with reported pain intensity, this analysis was also performed with demeaned pain ratings for each patient as a covariate. An F-test was used to test whether a difference between the conditions or pain ratings or a combination of these two contrasts had a non-zero effect.

Differences in perfusion between patients and controls during each condition separately were investigated using a between-group analysis with variables representing each group’s mean effect, the demeaned, age, sex and mean grey matter perfusion values for each participant entered as covariates of no interest.

A post hoc region of interest (ROI) analysis was performed to quantify the observed perfusion changes. Mean perfusion values were extracted from a sphere around the maximum voxel in activation clusters identified in the within-group analysis of x-axis flipped data in the patient group, so that the activation could be described as either ipsi- or contralateral.

To investigate the effect of group (patients versus controls) and condition (arm-up versus arm-down) a regression analysis was used with age, sex and the mean grey matter perfusion as covariates and with clustering for each subject. The percentage signal change between the conditions was also calculated in the patient group.

To investigate the expected multicollinearity between the pain rating and condition variables, a regression analysis was performed in the patient group first with and then without the condition variable; individual pain ratings, age, sex and mean grey matter perfusion values were entered as covariates and clustering was used to control for repeated measures.

## Results

### Participants characteristics

A total of 67 patients participated in the neuroimaging part of the CSAW trial. ASL data were not collected for five subjects, due to MRI contraindications, incidental findings, anxiety or excessive pain during the scanning session, unrelated to the evoked pain. Seven datasets were discarded due to technical reasons, such as incomplete brain coverage or missing calibration files. Two patients pressed the buzzer during the arm-up scan (scanning conditions are presented in Figure 1), which resulted in an acquisition of eight out of 10 epochs. For these two subjects, calibration files from the arm-down condition were used. These two datasets were included in the analysis as they did not differ from other datasets in terms of a perfusion pattern or mean and variance of grey matter perfusion. Twenty-four healthy controls were recruited; however, one could not be scanned due to MRI contraindications and one had incidental findings, while two datasets were discarded, due to excessive motion in one case and technical problems in the other. This left 55 patients and 20 control datasets for analysis.

Patient and control groups were matched in terms of age and sex. Mean age in the patient group was 51.5 years (Standard Deviation, SD 11.4, 95% Confidence iIterval, CI 48.4 to 54.6), 48.7 years in the healthy control group (SD 8.7, 95% CI 44.6 to 52.8) and the difference between the mean ages was not statistically significant (t = −0.998 p = 0.32). The male to female ratio was similar in both groups: 36% female in the patient group and 40% female in the control group (p = 0.79). There were also no differences in terms of which arm was raised during the arm-up scan: 39 out of 55 patients and 11 out of 20 controls raised their right arm during the arm-up condition (p = 0.27).

In the patient group, the median disease duration was 24 months (IQR 18 to 36), and 14 out of 55 patients were on medication that might have affected pain ratings. The median pain rating on the NRS for arm-down was 1 (IQR 0 to 4) and for arm-up 6 (IQR 5 to 8) (Figure 3) and the difference was statistically significant (Z = −5.8, p < 0.00005). The arm-down condition was painful for 51% of the patients, while the arm-up condition was painful for all of them, with neither being painful for healthy controls. Pain ratings associated with the shoulder pain during the arm-up condition were missing for two patients; therefore, analyses involving pain ratings were done in a total of 53 patients.

**Fig. 3.**
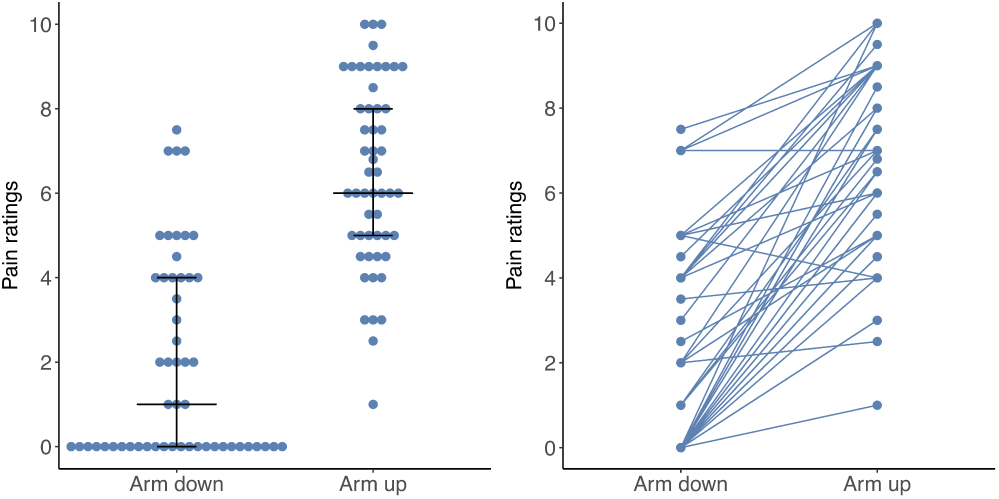
Ratings for pain intensity on a numerical rating scale (NRS) during the arm-down and arm-up condition in the patient group. Left panel: median with Interquartile range. Right panel: change in ratings between the arm-down and arm-up condition. Data are presented for N = 53 patients because arm-up pain ratings were missing for two patients.

### Mean grey matter perfusion per condition

The mean grey matter perfusion in the patient group during arm-down condition was 64.9 ml/100g/min (SD 10.9, 95% CI 62.0 to 67.9). During arm-up, this was 63.2 ml/100g/min (SD 12.2, 95% CI 59.9 to 66.5). In the control group, mean grey matter perfusion during arm-down was 62.3 ml/100g/min (SD 10.01, 95% CI 57.6 to 67.0). During arm-up, this was 61.2 ml/100g/min (SD 10.2, 95% CI 56.41 to 66.0).

The mean perfusion within the grey matter did not differ between patients and controls or the arm-up and arm-down condition and the interaction between the group and condition was also not significant. Both older age and male sex were associated with lower perfusion (Table 1).

**Table 1.** Linear regression of mean grey matter perfusion values (perfGM) adjusted for age, sex, condition (effect of arm-up versus arm-down), group (effect of being in the patient group versus control group) as well as the interaction between the condition and group (condition *#* group), clustered by subject (*regress perfGM age i.sex i.condition##i.group, cl(id)*). Abbreviations: *β* - regression coefficient, SE - standard error, t - t-statistic, p - p-value, CI - confidence interval

| perfGM | $\beta$ | SE | $t$ | $p$ | 95% CI |
| --- | --- | --- | --- | --- | --- |
| age | -0.34 | 0.08 | -4.08 | 0.00 | -0.51 to -0.17 |
| sex | -12.56 | 1.99 | -6.32 | 0.00 | -16.52 to -8.60 |
| cond. | -1.10 | 0.99 | -1.10 | 0.27 | -3.07 to 0.88 |
| group | 4.07 | 2.35 | 1.74 | 0.087 | -0.60 to 8.74 |
| condition#group | -0.60 | 1.27 | -0.48 | 0.64 | -3.13 to 1.92 |
| constant | 86.41 | 4.39 | 19.67 | 0.00 | 77.66 to 95.16 |

The arterial arrival time (AAT) values also did not differ for patients in comparison to controls or for arm-up in comparison to arm-down condition (Table 2).

**Table 2.** Linear regression of mean grey matter perfusion values (perfGM) adjusted for age, sex, condition (effect of arm-up versus arm-down), group (effect of being in the patient group versus control group), interaction between the condition and group (condition *#* group), clustered by subject (*regress arrivaltime age i.sex i.condition##i.group, cl(id)*). Abbreviations: *β* - regression coefficient, SE - standard error, t - t-statistic, p - p-value, CI - confidence interval

| AAT | $\beta$ | SE | t | p | 95% CI |
| --- | --- | --- | --- | --- | --- |
| age | 0.002 | 0.0007 | 2.88 | 0.005 | 0.0006 to 0.004 |
| sex | 0.11 | 0.014 | 7.90 | 0.000 | 0.08 to 0.14 |
| condition | -0.001 | 0.006 | -0.22 | 0.83 | -0.01 to 0.01 |
| group | -0.03 | 0.019 | -1.86 | 0.07 | -0.07 to 0.002 |
| condition#group | 0.0004 | 0.008 | 0.04 | 0.97 | -0.02 to 0.02 |
| constant | 1.006 | 0.04 | 27.16 | 0.00 | 0.93 to 1.08 |

### Perfusion differences between the arm-up and arm-down condition

The arm-up condition in comparison to arm-down was associated with increased perfusion in the contralateral primary motor and primary sensory, mid- and posterior cingulate cortex, and, bilaterally, in the operculum/insular cortex, putamen, inferior parietal lobule, thalamus, midbrain and the cerebellum (Figure 4 and Table 3). The interaction between the group and condition was not significant, except for a cluster in the cerebellum, which was likely to be an artefact caused by an incomplete coverage of cerebellum by the pCASL sequence.

**Table 3.**
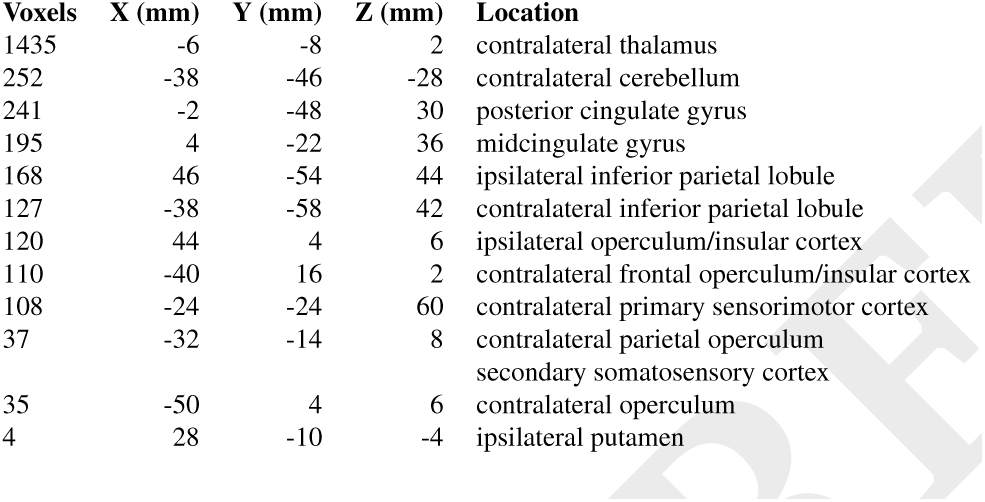
Coordinates of maximum p-values within each cluster for the main effect of condition presented in Figure 4

**Fig. 4.**
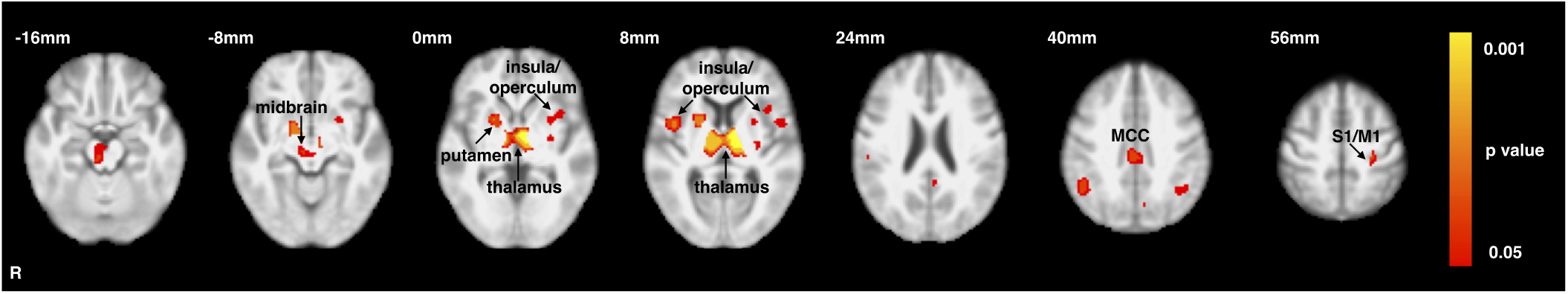
Differences between the arm-up and arm-down condition across the groups. Regression analysis in N = 55 patients and N = 20 control subjects. The main effect of condition across both groups, with the mean grey matter perfusion entered as a covariate of no interest. Perfusion images were flipped along the x-axis so that the right side of brain represented the ipsilateral side. The statistical image was Family-Wise Error-corrected using threshold-free cluster enhancement (TFCE) and thresholded to show voxels significant at p *<* 0.05. Abbreviations: S1 - primary somatosensory cortex, M1 - primary motor cortex.

### Perfusion differences between patients and healthy controls

Increased perfusion in the patient group in comparison with the control group was present, bilaterally, in the thalamus, midbrain, and the cerebellum, as well as in the ipsilateral temporal lobe (Figure 5 and Table 4).

**Table 4.** Coordinates of maximum p-values within each cluster for the main effect of condition presented in Figure 5

| Voxels | X (mm) | Y (mm) | Z (mm) | Location |
| --- | --- | --- | --- | --- |
| 5971 | 24 | -42 | -30 | cerebellum |
| 919 | 52 | -24 | -30 | ipsilateral inferior temporal gyrus |
| 96 | 28 | 46 | -10 | ipsilateral frontal pole |
| 10 | -30 | 46 | 36 | contralateral frontal pole |

**Fig. 5.**
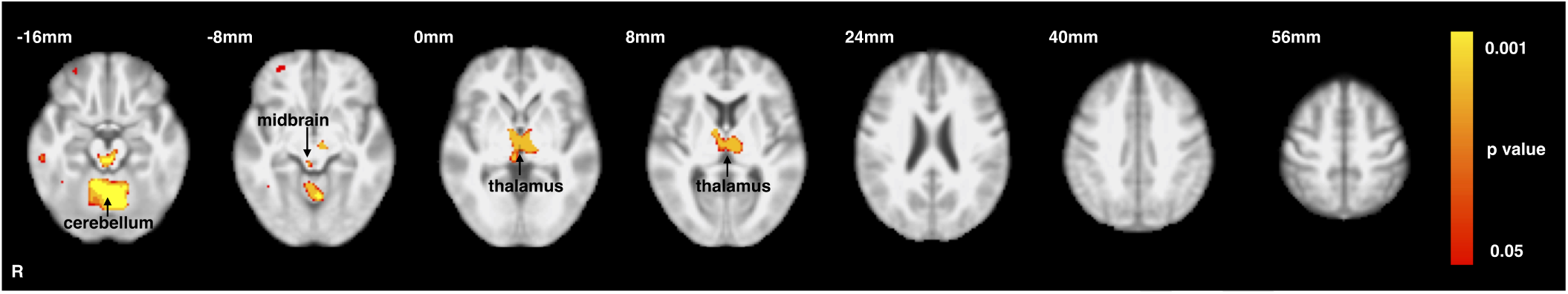
Differences between patients and controls across the conditions. Regression analysis in N = 55 patients and N = 20 control subjects. The main effect of group (patients > controls) across both conditions, with the age, sex, and mean grey matter perfusion entered as covariates of no interest. Perfusion images were flipped along the x-axis so that the right side of brain represented the ipsilateral side. The statistical image was Family-Wise Error-corrected using threshold-free cluster enhancement (TFCE). The results were thresholded to show voxels significant at p *<* 0.05.

### Post hoc analysis

In the patient group, there was an increase in perfusion between the arm-up and arm-down condition (patients arm-up > patients arm-down) in similar brain regions as the main effect of condition (Supplementary Figure S1). and Supplementary Table S1); there was no significant effect for the opposite contrast. In the control group, this analysis did not result in any significant clusters for the controls arm-up *>* controls arm-down contrast and the opposite contrast showed a small cluster in the occipital cortex, which was likely to be an artefact because of inconsistent brain coverage across the scans.

When individual pain ratings were included in the within-group analysis, neither the contrast representing a difference between the conditions nor the contrast representing individual pain ratings resulted in a significant effect. However, an F-test of these two contrasts resulted in a significant cluster, which extended from the thalamus, through the midbrain to the cerebellum, with the peak voxel in the left cerebellum (maximum p-value coordinates in mm: −34, −62, −36) (Supplementary Figure S2)

A between-group analysis of the perfusion images during arm-down condition (patients arm-down > controls arm-down) did not result in significant differences. The same analysis for the arm-up condition (patients arm-up > controls arm-up), showed an increased perfusion in a cluster encompassing the thalamus, midbrain and the cerebellum; with additional clusters in the temporal lobes and orbitofrontal cortex (Supplementary Figure S3) and Supplementary Table S2)). There were no significant results for the opposite contrast (controls arm-up > patients arm-up).

### Post hoc ROI analysis

A post hoc ROI analysis was performed within three regions, which showed significant differences in analysis between arm-up and arm-down condition in the patient group presented in Figure 1. The mean perfusion values were extracted from spherical ROIs with 5mm radius and peak in the contralateral primary somatosensory cortex (coordinates in mm: −26, −26, 56), the contralateral operculum (coordinates in mm: −50, 4, 8) and the contralateral thalamus (coordinates in mm: −10, −8, 6).

A regression analysis of the effect of the group and condition (Table 5), demonstrated a significant effect of the condition but not the group, both, in the contralateral primary sensory cortex and in the contralateral operculum. For the contralateral thalamus, both effects were statistically significant. In all three ROIs, the effect of the mean grey matter perfusion was highly significant (p*<*0.0005).

**Table 5.** Linear regression of mean perfusion values within each region of interest (ROI) adjusted for age, sex, mean grey matter perfusion (perfGM), condition, group(effect of being in the patient group versus being in the control group) and clustered by subject (id). Data for the arm-up and arm-down condition in the patient and control group (number of observations = 150) (*regress ROI age i.sex i.condition i.group, cl(id)*). Abbreviations: S1 – primary somatosensory cortex, *β* – regression coefficient, SE – standard error, t – t-statistic, p - p-value, CI – confidence interval.

| S1 | $\beta$ | SE | t | p | 95% CI |
| --- | --- | --- | --- | --- | --- |
| age | 0.12 | 0.11 | 1.06 | 0.29 | -0.11 to 0.35 |
| sex | -0.89 | 2.40 | -0.37 | 0.71 | -5.67 to 3.90 |
| perfGM | 0.83 | 0.10 | 8.16 | 0.00 | 0.63 to 1.04 |
| condition | 3.86 | 0.72 | 5.35 | 0.00 | 2.42 to 5.30 |
| group | 1.45 | 2.48 | 0.58 | 0.56 | -3.49 to 6.38 |
| constant | -19.33 | 10.22 | -1.89 | 0.06 | -39.70 to 1.04 |

| Operculum | $\beta$ | SE | t | p | 95% CI |
| --- | --- | --- | --- | --- | --- |
| age | -0.02 | 0.10 | -0.19 | 0.85 | -0.22 to 0.18 |
| sex | 0.58 | 1.78 | 0.33 | 0.74 | -2.95 to 4.12 |
| perfGM | 0.65 | 0.07 | 8.90 | 0.00 | 0.51 to 0.80 |
| condition | 3.30 | 0.73 | 4.50 | 0.00 | 1.84 to 4.77 |
| group | 2.22 | 1.45 | 1.53 | 0.13 | -0.66 to 5.10 |
| constant | 18.03 | 8.31 | 2.17 | 0.03 | 1.46 to 34.59 |

| Thalamus | $\beta$ | SE | t | p | 95% CI |
| --- | --- | --- | --- | --- | --- |
| age | 0.07 | 0.09 | 0.80 | 0.43 | -0.11 to 0.25 |
| sex | -3.40 | 1.42 | -2.39 | 0.02 | -6.24 to -0.57 |
| perfGM | 0.71 | 0.07 | 10.27 | 0.00 | 0.57 to 0.85 |
| condition | 2.98 | 0.56 | 5.28 | 0.00 | 1.85 to 4.10 |
| group | 4.17 | 1.71 | 2.43 | 0.02 | 0.75 to 7.58 |
| constant | 1.50 | 7.58 | 0.20 | 0.84 | -13.61 to 16.61 |

Patients rated the arm-up condition as painful and arm-down condition as slightly painful or not painful, which caused multicollinearity between the condition and pain ratings variable. When both variables were entered in the regression model, ROI perfusion values in the contralateral primary somatosensory cortex did not correlate with individual pain ratings but the effect of condition was significant. For the con-tralateral operculum and contralateral thalamus, neither effect was significant (Table 6). When the condition variable was removed from the model, the perfusion correlated with pain ratings in the operculum and thalamus, but not in the primary somatosensory cortex (Table 7).

**Table 6.**
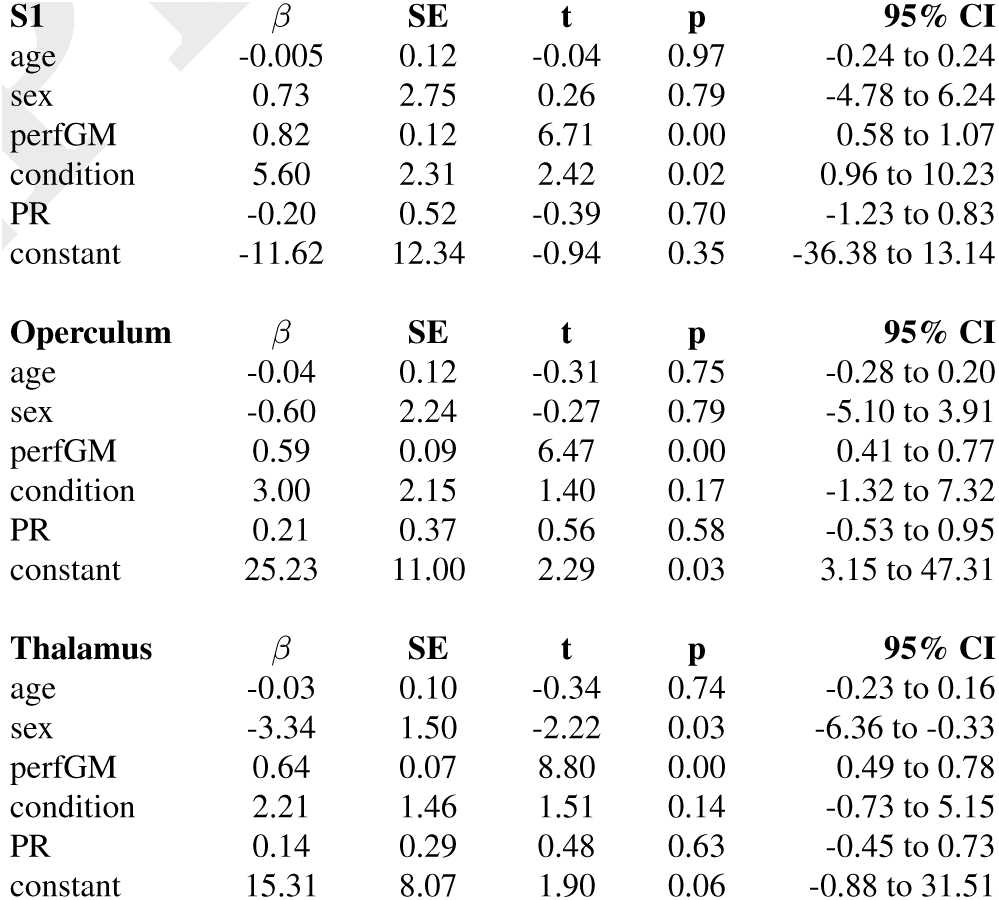
Linear regression of perfusion changes within each region of interest (ROI) adjusted for age, sex, the mean grey matter perfusion (perfGM), the condition, pain ratings (PR), and clustered by subject (id). Only data for the arm-up and arm-down condition in the patient group were analysed. Pain ratings for the arm-up condition were missing for two patients; therefore, data of 53 patients were included in the analysis (number of observations = 106) (*regress ROI condition PR perfGM age i.sex, cl(id)*). Abbreviations: S1 – primary somatosensory cortex, *β* – regression coefficient, SE – standard error, t – t-statistic, CI – confidence interval.

**Table 7.**
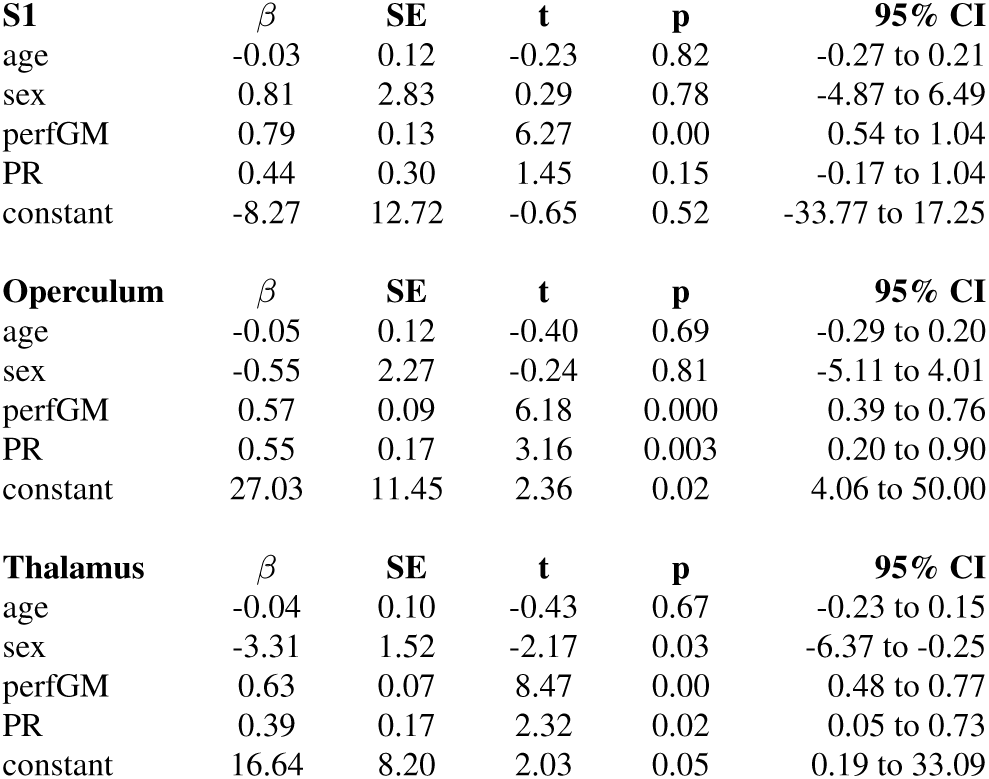
Linear regression of perfusion changes within each region of interest (ROI) adjusted for age, sex, and the mean grey matter perfusion, pain ratings but without the condition variable, clustered by subject (id). Only data for the arm-up and arm-down condition in the patient group were analysed. Pain ratings for the arm-up condition were missing for two patients; therefore, data of 53 patients were included in the analysis (number of observations = 106)(*regress ROI age sex perfGM PR, cl(id)*). Abbreviations: S1 – primary somatosensory cortex, *β* – regression coefficient, SE – standard error, t – t-statistic, CI – confidence interval.

In the patient group only, changes in ROI perfusion between the arm-up and arm-down condition were small. In the contralateral primary somatosensory cortex ROI, the increase was from 41.4ml/100g/min (SD 12.2) to 44.7 ml/100g/min (SD 14.5) (t = −3.4, p = 0.0014), corresponding to a difference of 8%; for the ROI in the contralateral operculum, the perfusion change was from 61.6 ml/100g/min (SD 9.6) to 64.56 ml/100g/min (SD 11.6) (t = 2.9 p = 0.006), which is 5%; for the ROI in the contralateral thalamus, the difference was from 53.3 ml/100g/min (SD 10.08) to 55.1 ml/100g/min (SD 11.3) (t = 1.99, p = 0.052), which is 3%.

## Discussion

The raising of the affected arm resulted in increased pain ratings and increased brain perfusion in the patient group. The arm-up condition was associated with a significant increase in perfusion in cortical and subcortical brain regions involved in sensory processing and movement integration. Perfusion differences between patients and controls were present bilaterally in the thalamus, midbrain, and cerebellum. The effect of interaction between the condition and group was significant only in the cerebellum; however, this effect is likely to be an artefact. In this study, the effect of condition could not be distinguished from the effect of pain rating because of multicollinearity between these two variables. When the condition variable was removed from the model, there was a significant association between pain ratings and perfusion in the contralateral operculum and thalamus.

The main strength of this study was that it involved a clinically-meaningful stimulus, which evoked perfusion changes related to the ongoing musculoskeletal pain that patients experience as a result of their condition. Secondly, it used a whole-brain analysis rather than pre-specified ROIs. Moreover, it had the advantage of using both within- and between-subject analyses, which helped with the interpretation of the observed results. Also, the sample size was two to three times larger than the previous ASL patient studies (3, 5, 18). Finally, the optimised analysis pipeline resulted in a robust registration of individual perfusion images to the group template, which was superior to standard registration.

This study has also some limitations. Firstly, ASL is less sensitive to changes than BOLD; therefore, it is likely that, in our study, the sample size was not large enough to show the interaction between the group and condition or to show a correlation between perfusion and pain ratings. An earlier study (19) demonstrated that 10 to 15 healthy controls would be required to observe 15% changes in a within-subject model. In our study, there were 55 patients but the magnitude of signal change between conditions in the patient group was around 3-8%. Data from older subjects, especially patients, require a larger sample due to variance in the perfusion data related to physiological factors and clinical heterogeneity. A recent paper suggested that at least 37 older participants per group would be required to detect a 10% perfusion difference using permutation-based algorithms (20). Similarly to Wasan and colleagues (3), we did not observe a significant correlation between changes in pain ratings and changes in perfusion. They interpreted this as an indication that either the changes in pain ratings were too small (they expected to see a change in perfusion related to pain only when pain ratings increased > 30%) or that perfusion change was not a marker of pain severity (3). However, the correlation between clinical pain ratings and perfusion in several brain regions, including in the ipsilateral insular cortex and operculum, putamen, the amygdala/hippocampus region, and brainstem, was reported by another study (4). In our study, the lack of significant correlations between changes in pain ratings and perfusion was most likely caused by small sample size, as well as by multicollinearity between the condition and pain ratings. Most of the difference between pain ratings during the arm-up and arm-down conditions was accounted for by the condition, leaving very little variance that could be explained by the covariate representing the reported pain ratings. When the condition was removed from the model, perfusion in the contralateral thalamus and operculum correlated with pain ratings.

Another limitation was that the order of arm-down and arm-up scans could not be randomised due to the possibility of the evoked pain persisting beyond the duration of the scan; therefore, the order effect may be a confounder. However, we would not expect a substantial carry-over effect from patients lying in the scanner with arms beside the body. It is more likely they were more restless during the more painful and uncomfortable arm-up scan, especially as it was performed at the end of a long scanning session. This effect might have been exacerbated in patients who experienced pain during the arm-down condition.

In this study, differences between the arm-up and arm-down condition were observed in the structures encoding the sensory-discriminatory dimension of pain, including pain intensity (21–24). A post hoc analysis demonstrated that differences between the arm-up and arm-down were significant in the patient but not in the control group; therefore, they might be interpreted as related to pain rather than arm repositioning. A similar increase in perfusion in the primary and secondary somatosensory cortex, mid-insula, amygdala, hippocampus, and midbrain was reported in an earlier ASL study on osteoarthritis (18). Increased activity in the primary somatosensory cortex, anterior cingulate, insula, midbrain, thalamus and cerebellum has been also described during capsaicin-induced sensitisation in healthy volunteers (25). Therefore, observed changes may be interpreted as altered pain processing related to central sensitisation. However, as the effect was present in the primary somatosensory cortex as well as in the thalamus and putamen, it may be also interpreted as reflecting maladaptive changes in the network integrating movement and sensory information (26–28).

The interaction term for group and condition demonstrated significant increase in perfusion in the cerebellum; however, this result might an artefact because the pCASL sequence did not cover the whole cerebellum. The study was most likely underpowered to detect a significant effect of interactions.

Differences between patients and controls were present mainly in the thalamus, which is the main relay centre for the pathways transmitting nociceptive information and has been reported to be involved in chronic pain processing (23, 29, 30). There was also a cluster in the midbrain, possibly associated with modulation of thalamic transmission (31). Higher levels of BOLD activation in patients than controls in the brainstem, thalamus and cerebellum have previously been described in response to pressure pain in fibromyalgia (32). A recent ASL study in fibromyalgia patients reported that perfusion in the putamen correlated negatively with pain disability, whereas pain intensity correlated with perfusion in the cerebellum (28).

## Conclusions

In conclusion, arm repositioning may be used as a paradigm for evoked ongoing musculoskeletal pain in patients with shoulder pain. In this study, raising the affected arm resulted in perfusion changes in cortical and subcortical regions. This effect could be interpreted as a result of central sensitisation or changes in the network integrating movement and sensory information. The main drawback was low sensitivity of ASL, which precluded detection of small perfusion changes. Registration between the ASL and the structural space, which was challenging due to the low between-tissue contrast inherent to ASL, was improved by optimising the pre-processing steps. ASL is useful to study musculoskeletal pain in patient’s population.

## ACKNOWLEDGEMENTS

The authors would like to thank the following organisations who provided funding for this work: EPSRC UK (EP/P012361/1), NIHR Oxford Musculoskeletal Biomedical Research Unit and the Royal Academy of Engineering. The funders had no role in study design, data collection and analysis, decision to publish or preparation of the manuscript. The authors would like to thank the SITU team who ran the main CSAW trial, Dr Anderson Winkler who provided important feedback for statistical analyses and Dr Maria Kuzma-Kuznarska (http://mybioscience.org/) for creating the drawing in Figure 1.

## AUTHOR CONTRIBUTIONS

KAW designed the study, acquired and analysed the data. They also drafted the manuscript and acted as the guarantor of the study. KAW, DPB, MAC, MJ, TWO, MAW and AJC contributed to the analysis and writing of the manuscript, also critically revised it for important intellectual content. All authors agree with the manuscript’s results and conclusions.

## DISCLAIMER

The authors declare that there is no conflict of interest.

## A. Appendix

**Fig. S1.**
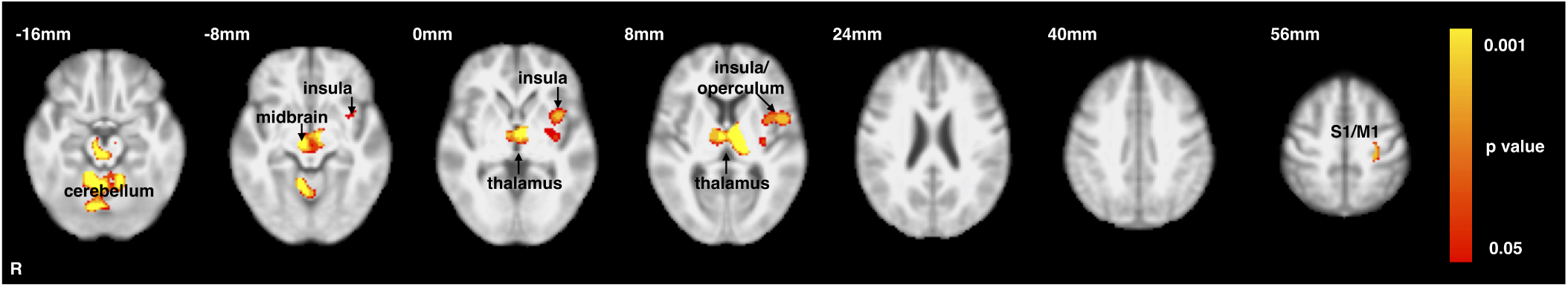
Differences between the arm-up and arm-down condition in the patient group. Within-group analysis in N = 55 patients, comparing patients arm-up > patients arm-down. Perfusion images were flipped along the x-axis so that the right side of brain was the ipsilateral side. The statistical image was Family-Wise Error-corrected using threshold-free cluster enhancement (TFCE) and thresholded to show voxels significant at p *<* 0.05. Abbreviations: S1/M1 – primary sensorimotor cortex.

**Table S1.**
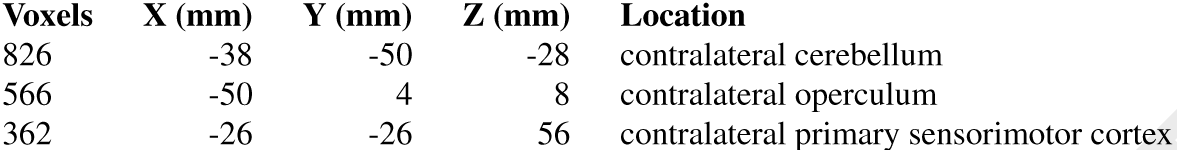
Coordinates of maximum p-values within each cluster for the main effect of condition presented in Figure S1

| Voxels | X (mm) | Y (mm) | Z (mm) | Location |
| --- | --- | --- | --- | --- |
| 826 | -38 | -50 | -28 | contralateral cerebellum |
| 566 | -50 | 4 | 8 | contralateral operculum |
| 362 | -26 | -26 | 56 | contralateral primary sensorimotor cortex |

**Fig. S2.**
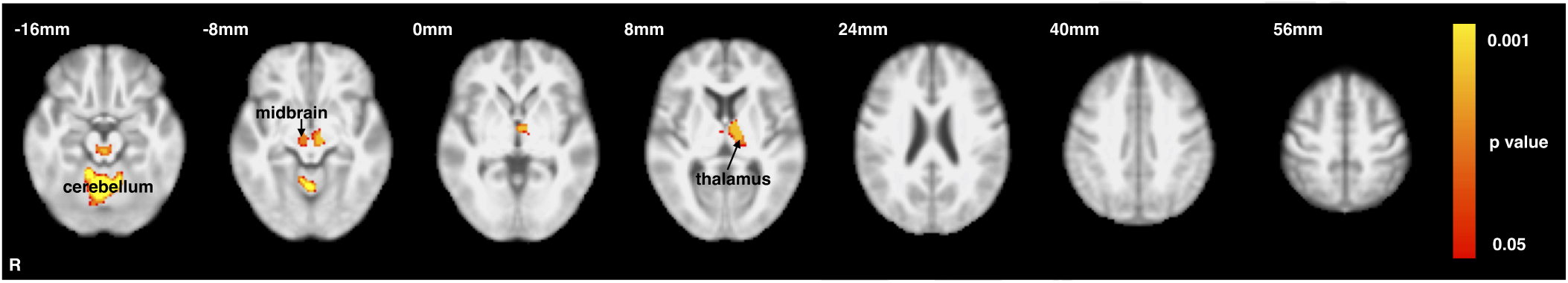
F-test for the within-group differences contrast and pain ratings contrast in the patient group. F-test results for the patients arm-up *>* patients arm-down contrast and the individual pain ratings contrast in a within-group analysis with the mean grey matter perfusion as a covariate of no interest. Analysis was performed in N = 53 patients because arm-up pain ratings were missing for two patients. Perfusion images were flipped along the x-axis so that the right side of brain represented the ipsilateral side. The statistical image was Family-Wise Error-corrected using threshold-free cluster enhancement (TFCE) and thresholded to show voxels significant at p *<* 0.05.

**Fig. S3.**
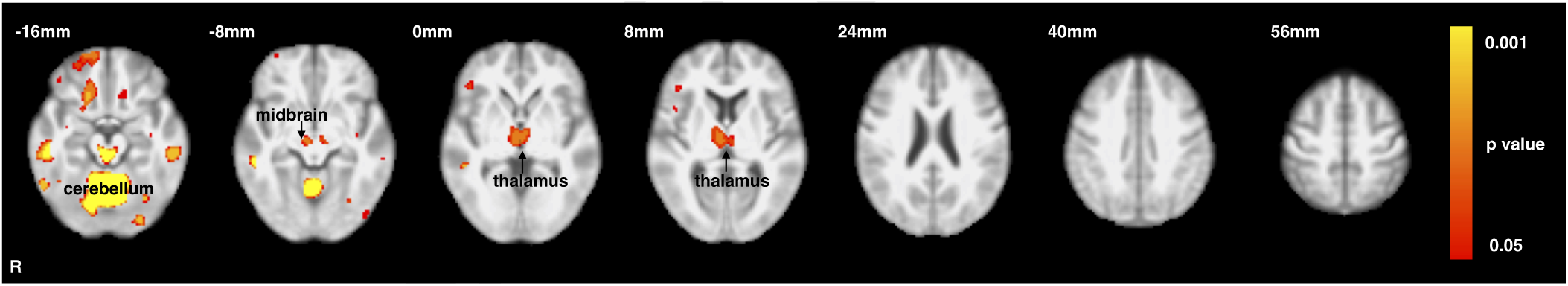
Between-group differences for the arm-up condition. Between-group analysis in N = 55 patients and N = 20 control subjects during the arm-up condition (patients arm-up *>* controls arm-up), with age, sex, and the mean grey matter perfusion as covariates of no interest; flipped along the x-axis so that the right side of brain represented the ipsilateral side. The statistical image was Family Wise Error-corrected using threshold-free cluster enhancement (TFCE). The results were thresholded to show voxels significant at p *<* 0.05.

**Table S2.** Coordinates of maximum p-values within each cluster for the main effect of condition presented in Figure S3.

| Voxels | X (mm) | Y (mm) | Z (mm) | Location |
| --- | --- | --- | --- | --- |
| 8718 | 30 | -42 | -38 | cerebellum |
| 2039 | 54 | -26 | -30 | ipsilateral inferior temporal gyrus |
| 1291 | -44 | -26 | -26 | contralateral inferior temporal gyrus |
| 638 | 18 | 26 | -18 | ipsilateral orbitofrontal cortex |
| 60 | -12 | 26 | -20 | contralateral fronto-orbital cortex |
| 7 | -30 | 46 | -12 | contralateral frontal pole |
| 6 | -46 | -78 | -12 | contralateral lateral occipital cortex |
| 5 | -30 | -68 | -10 | contralateral fusiform gyrus |
| 3 | -36 | -14 | -14 | contralateral hippocampus |

